# Partial loss of colonic primary cilia promotes inflammation and carcinogenesis

**DOI:** 10.1101/2019.12.20.871772

**Authors:** Ruizhi Tang, Conception Paul, Rossano Lattanzio, Thibaut Eguether, Hulya Tulari, Julie Bremond, Chloé Maurizy, Sophie Poupeau, Andrei Turtoi, Magali Svrcek, Philippe Seksik, Vincent Castronovo, Philippe Delvenne, Bénédicte Lemmers, Carsten Janke, Valérie Pinet, Michael Hahne

## Abstract

Primary cilia (PC) are important signaling hubs in cells and their deregulation has been associated with various diseases including cancer. Here we explored the role of PC in colorectal cancer (CRC) and colitis. In the colon we found PC to be mostly present on different subtypes of fibroblasts. Colons of mice exposed to either chemically induced colitis-associated colon carcinogenesis (CAC) or dextran sodium sulfate (DSS)-induced colitis had decreased numbers of PC. We employed conditional knock-out strains for the PC essential genes, *Kif3A* and *Ift88*, to generate mice with reduced numbers of PC on colonic fibroblasts. These mice showed an increased susceptibility in the CAC model as well as in DSS-induced colitis. Colons from DSS-treated mice with PC-deficiency on fibroblasts displayed an elevated production of the pro-inflammatory cytokine IL-6 and colonic epithelial cells had diminished levels of HES-1, a key transcription factor of Notch signaling. Notably, an analysis of PC presence on biopsies of patients with ulcerative colitis as well as CRC patients revealed decreased numbers of PC on colonic fibroblasts in pathological versus surrounding normal tissue. Taken together, we provide evidence that a decrease in colonic PC numbers promotes colitis and CRC.

**Graphical Abstract:** 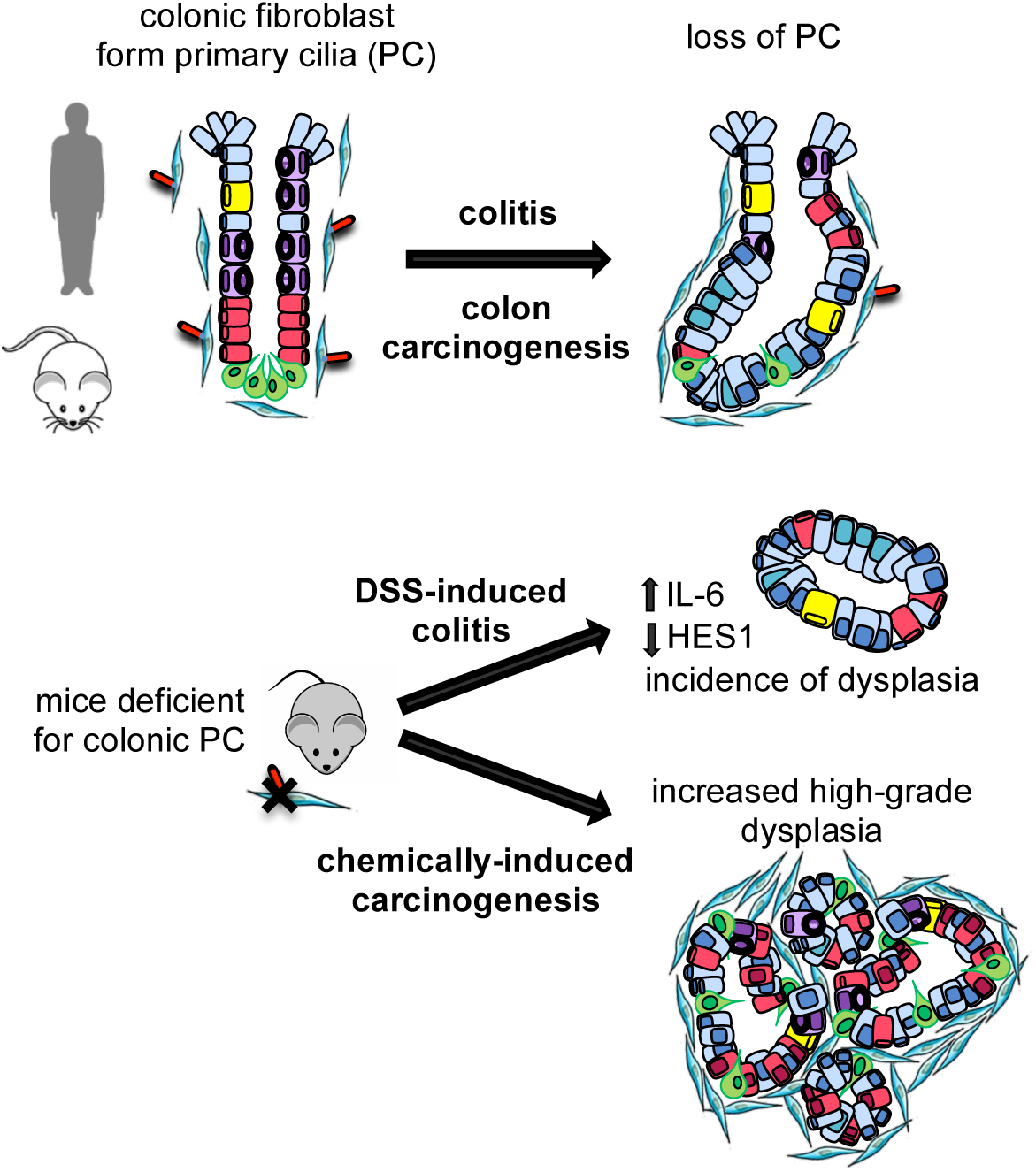

## Introduction

Colorectal cancer (CRC) is the third most common and deadly cancer worldwide (Malvezzi *et al*, 2018). Due to its asymptomatic nature, CRC is frequently diagnosed only after the cancer has spread through the colon or rectal wall (stage II), to the lymph nodes (stage III) and/or to distant organs (stage IV), e.g. liver, lung and peritoneum. CRC is a molecularly highly heterogeneous disease and there is an urgent need to map the heterogeneity of CRC, to identify novel therapeutic intervention points as well as novel prognostic and predictive biomarkers that pre-select stage II and III CRC patients who will not benefit from standard-of-care chemotherapy, re-directing them towards novel, targeted treatment options. In addition, clinical studies demonstrate that innate and/or acquired tumor resistance remains a core problem in the management of stage IV CRC patients, further highlighting the need to develop biomarker-guided therapeutic approaches for treatment of those patients.

A recent meta-analysis on the different reported sub-classifications defined the existence of at least 4 distinct gene expression–based CRC subtypes, called Consensus Molecular Subtypes (CMS) 1-4, each displaying unique biology and gene expression pattern (Guinney *et al*, 2015). Among those CMS4 cancers have the worst prognosis and are characterized by a stromal and inflammatory gene signature (Becht *et al*, 2016). The importance of inflammation during tumorigenesis is well illustrated by the fact that tissues with chronic inflammation have an increased risk to develop tumors (Grivennikov et al, 2009). Hence, patients with inflammatory bowel disease have a higher probability to develop CRC (Beaugerie et al, 2017).

A master regulator of signaling pathways are primary cilia (PC), which are supposed to extrude most mammalian cells and act as sensory antennae (Gerdes *et al*, 2009). PC contain a scaffold of nine microtubule (MT) doublets forming a cylinder-like arrangement in the plasma membrane by a basal body that derives from a centriole. PC are involved in a variety of diseases commonly referred to as ciliopathies including cystic kidney diseases (Ko & Park, 2013), but they are also involved in tumor formation (Han *et al*, 2009)(Wong *et al*, 2009). In fact, a picture is emerging that there is an important inter- and intra-tumoral variation of PC numbers (Liu *et al*, 2018)(Eguether & Hahne, 2018). PC properties can be modulated by post-translational modifications (PTMs) of tubulin including acetylation and glycylation (Wloga *et al*, 2017). For example, it has been shown that blocking of deacetylation induces cilia restoration and decreases tumor growth of cholangiocarcinoma (Gradilone *et al*, 2013). We previously described an unexpected role of the tubulin glycylase TTLL3 in the regulation of colon tumorigenesis (Rocha *et al*, 2014). Specifically, we discovered that TTLL3 is the only glycylase expressed in the colon and that the absence of TTLL3 leads to decreased numbers of PC in colonic crypts (Rocha *et al*, 2014). When exposed to chemically induced colon carcinogenesis, *Ttll3*^-/-^ mice are more susceptible to tumor formation (Rocha *et al*, 2014). Importantly, TTLL3 expression levels were significantly downregulated in human primary colorectal carcinomas and metastases as compared to healthy colon tissue, strongly suggesting a pivotal role of TTLL3 in colorectal cancer. All together, these findings revealed a first direct link between the downregulation of PC and colon carcinogenesis.

In this study, we directly investigated the role of PC in colon carcinogenesis by employing relevant mouse models and analysis of patient biopsies.

## Results

### Colonic fibroblasts form primary cilia

To characterize the presence of PC in the colon we employed a previously described protocol that allows the detection of PC on paraffin-embedded tissues (Hassounah *et al*, 2013). Co-staining was performed combining the established PC marker Arl13b (Caspary *et al*, 2007) with markers for epithelial or stromal cells including E-cadherin, vimentin, alpha-smooth muscle actin (α-SMA) and CD140a (platelet-derived growth factor receptor alpha). Staining for vimentin and α-SMA allows to distinguish fibroblasts from myofibroblasts, whereas CD140a characterizes a recently described vimentin^low^ α-SMA^negative^ subpopulation of fibroblasts present on the upper parts of the crypts (Roulis & Flavell, 2016)(Kurahashi *et al*, 2013) (Figures 1 B-D). While only few epithelial cells displayed detectable PC (Figure 1A and supplementary Figure S1), PC were mostly found on vimentin^positive^ or CD140a^positive^ stromal cells in the lamina propia (Figures 1B-D), but also on α-SMA^+^ myofibroblasts (Figure 1D, E).

**Figure 1.**
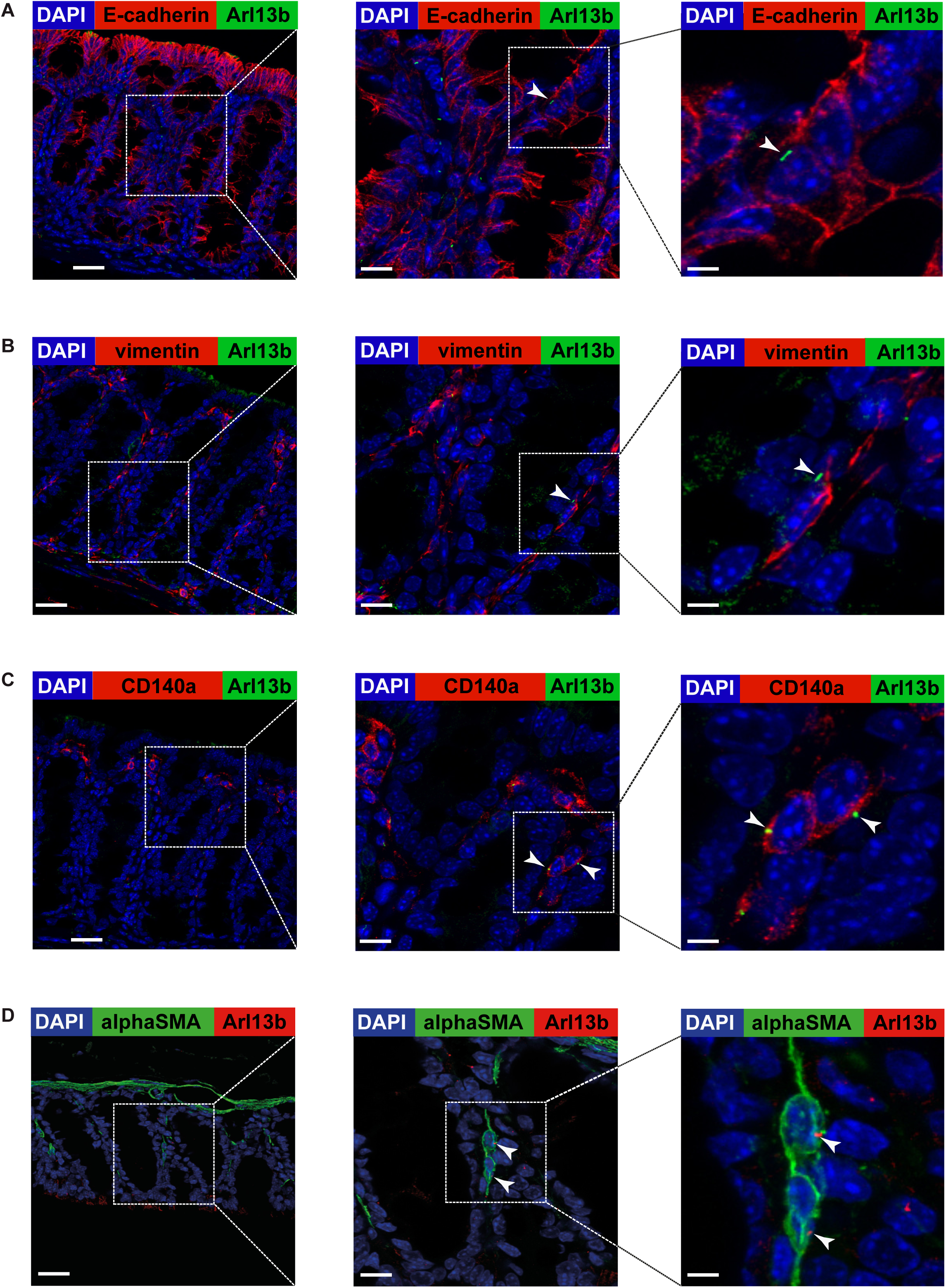
Primary cilia in the colon are mostly present on stromal cells. (A-D) PC were identified by immunostaining for Arl13b (in green except in (D); arrowheads in the panel on the right-hand side), and in parallel by identifying epithelial cells with E-cadherin(A), fibroblasts with vimentin (B) and CD140a (C), labeling (all in red), and myofibroblasts by α-SMA stain (D) in green). Nuclei were stained with DAPI (in blue). Scale bars represent 30, 10 and 3 µm, respectively (from the left to the right).

### Progressive loss of colonic primary cilia during colon carcinogenesis

To determine whether decreased numbers of PC promote colon carcinogenesis we first analyzed the presence of PC in a mouse model that mimics colitis-associated colon carcinogenesis (CAC) (Tanaka *et al*, 2003)(Suzuki *et al*, 2004). This model depends on the administration of the mutagen azoxymethane (AOM) and the subsequent induction of inflammation with dextran sodium sulfate (DSS) (Figure 2A). DSS is toxic to mucosal epithelial cells in the colon, and the ensuing breakdown of the mucosal barrier leads to inflammation. Colonic crypts displayed a lower number of PC on vimentin^positive^ cells in tumor lesions as compared to adjacent normal tissue (Figures 2B,C). Remarkably, areas with high-grade dysplasia showed significant lower numbers of PC than those with low-grade dysplasia. Taken together, CAC is associated with a down-regulation of PC, confirming our initial hypothesis.

**Figure 2.**
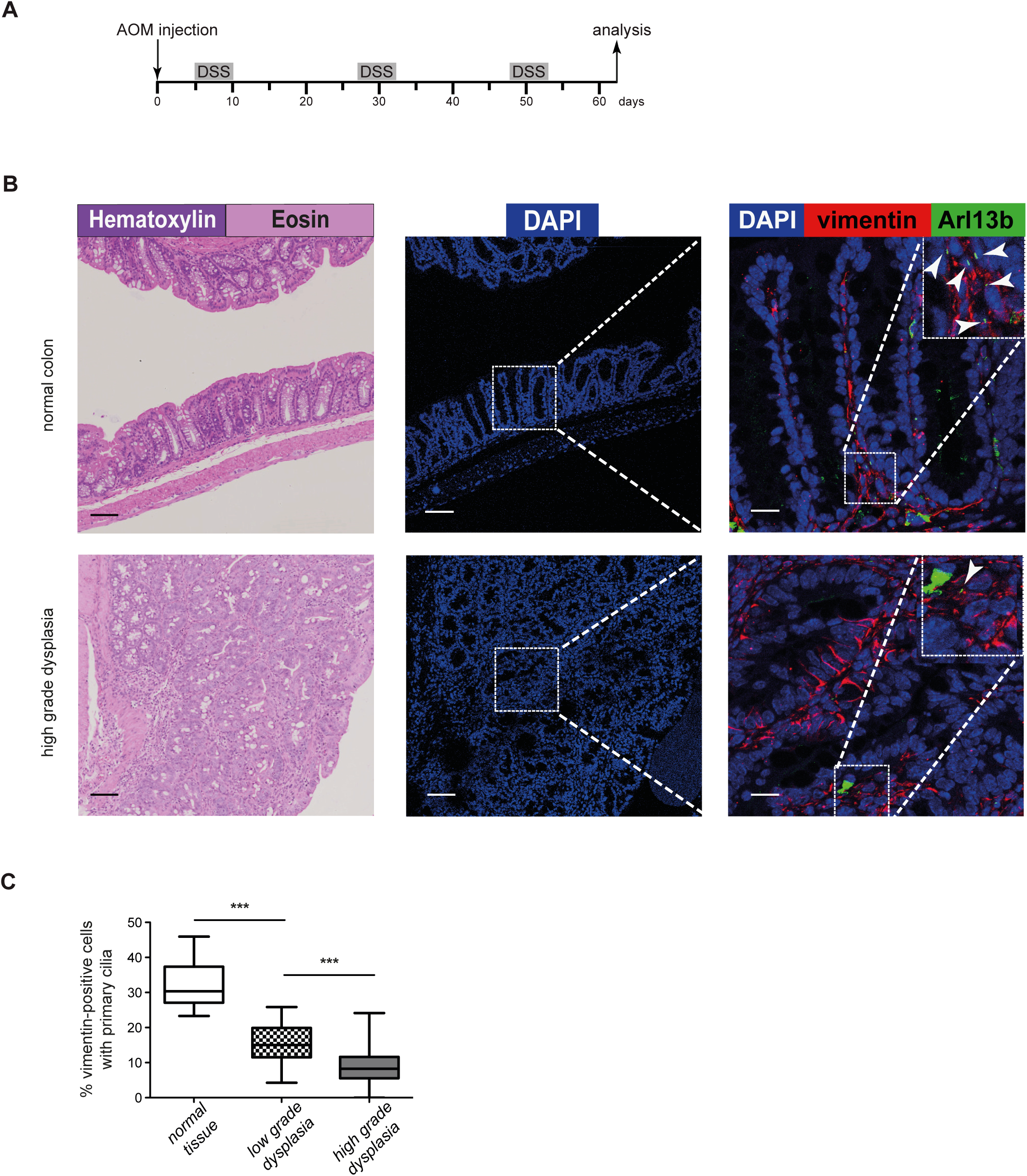
Number of colonic primary cilia decreases during colon carcinogenesis. (A) Experimental timeline for azoxymethane (AOM)/dextran sodium sulfate (DSS)-induced CAC. (B) HE and corresponding confocal images (maximum projection) of paraffin-sections from colons of AOM/DSS treated mice as indicated in (A). A representative example for a normal region and an area with high grade dysplasia are shown. Sections were stained with DAPI (blue) and an anti-Arl13b antibody (green) that specifically labels PC (arrowheads). Colonic fibroblasts were revealed with an anti-vimentin antibody (red). Scale bars represent 100 µm in the panels in the middle and on the left side and 20 µm in the panel on the right side. The panel on the right side includes 2,5x blow-ups. (C) Box-and-whisker plots showing quantitative analysis of PC expression on vimentin^+^ cells in normal colon, and colon with low and high grade dysplasia in mice exposed to the AOM/DSS protocol (analyzing at least 12 fields depicted from 4 different mice as illustrated in middle panel of (B)). Significance values were determined by two-tailed unpaired t-tests (****p<10^−4^).

### Decreased numbers of primary cilia in colonic fibroblasts promote CAC

To establish a causal relationship of PC on colonic fibroblasts and the development of CRC, we employed a mouse strain carrying conditional knockout (KO) alleles for the kinesin family member 3A (*Kif3a*^*flx/flx*^) (Marszalek *et al*, 1999), a protein essential for cilia formation. Tissue-specific deletion of this gene in mesenchymal cells, including intestinal fibroblasts, was achieved in combination with ColVI-cre transgenic mice, expressing Cre DNA-recombinase under the control of a collagenase VI promoter (Armaka *et al*, 2008)(Koliaraki *et al*, 2015)(Roulis & Flavell, 2016). ColVIcre*-Kif3a*^*flx/flx*^ mice were fertile, born at the expected mendelian ratio and displayed no overt intestinal phenotype (Figure 3A,B). *Kif3a*-deletion was indeed only detectable in intestinal fibroblasts, but not epithelial cells (Supplementary Figure S2A) and resulted in significantly less PC in the colon of ColVIcre-*Kif3a*^*flx/flx*^ mice, although the depletion of PC on vimentin^positive^ and CD140α^positive^ colonic fibroblasts was only partial (30-40%, Figures 3C,D). This is in line with a previous report that the ColVI promoter is active only in a part of vimentin^positive^ or CD140a^positive^ colonic fibroblasts (Koliaraki *et al*, 2015). Moreover, biological and clinical differences for different segments of the colon have been described (Minoo *et al*, 2010), but the decrease of PC numbers ColVIcre-*Kif3a*^*flx/flx*^ mice is consistent across distal, transversal and proximal regions of the colon (Supplementary Figure 2B).

**Figure 3.**
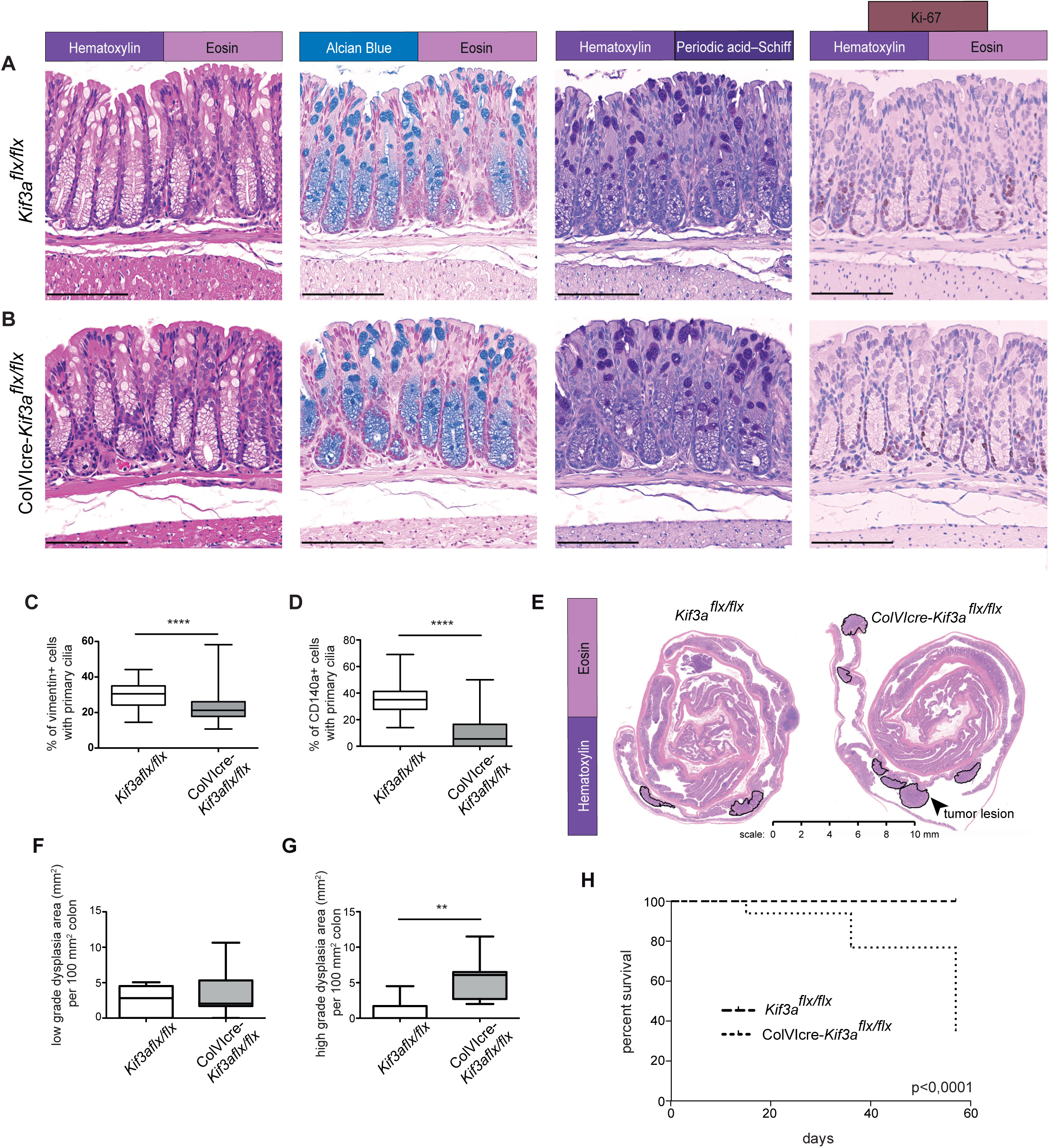
Decreased number of primary cilia in colonic fibroblasts promotes CAC. (A,B) No alterations in the architecture between colons of *Kif3a*^*flx/flx*^ (A) and ColVIcre-*Kif3a*^*flx/flx*^ (B) mice are detectable. Representative images for stains with hematoxylin eosin, alcian blue and periodic acid-Schiff (PAS), as well as immunostain for the proliferation marker Ki-67 are shown. Alcian blue stains negatively charged mucins, whereas PAS recognizes neutral mucins as well. Scale bars represent 100 µm (C,D) Quantitative analysis of PC expression on vimentin^+^ (C) and CD140a^+^ (D) cells in colons of *Kif3a*^*flx/flx*^ *and* ColVIcre-*Kif3a*^*flx/flx*^ mice as described in Figure 2C. (E,F,G) CAC was induced in control (*n* = 7) and ColVIcre-*Kif3a*^*flx/flx*^ (*n* = 7) female mice according to the timeline shown in Figure 2A. (E) Representative images of hematoxylin and eosin staining of paraffin-embedded colon sections prepared as “Swiss roll”. High dysplasia lesions are delimited by black lines. (F,G) Box-and-whisker plots depicting area of low (F) and high grade (G) dysplasia per mouse with a line indicating the median values. (H) Kaplan-Meier survival curves of AOM/DSS-treated male *Kif3a*^*flx/flx*^ (n=13) and ColVIcre-*Kif3a*^*flx/flx*^ (n=11) mice. CAC was induced as described for Figure 2A. (C,D,G) **p<0.01, ****p<10^−4^ by two-tailed unpaired t-test.

We next tested whether the reduced number of PC in the colon of ColVIcre-*Kif3a*^*flx/flx*^ mice alters their susceptibility to AOM/DSS-induced carcinogenesis. Female ColVIcre-*Kif3a*^*flx/flx*^ mice displayed an increased incidence of dysplasia, in particular, a higher number of high-grade dysplasia as compared to *Kif3a*^*flx/flx*^ mice (Figures 3E,F,G). Male mice are known to be more susceptible to colon carcinogenesis than females (Lee *et al*, 2016) and indeed, following the AOM/DSS treatment we observed a survival rate of about 35% of ColVIcre*-Kif3a*^*flx/flx*^ males in contrast to 100% of controls (Figure 3H). Thus, PC deficiency in colonic fibroblasts enhances the sensitivity of mice to colitis-associated carcinogenesis.

### Mice deficient of primary cilia are more susceptible to DSS-induced colitis

AOM/DSS treated male ColVIcre-*Kif3a*^*flx/flx*^ mice displayed a significantly lower weight during the first and third treatment of DSS (Supplementary Figure S3). Moreover, the ColVIcre-*Kif3a*^*flx/flx*^ mice that died during the AOM/DSS protocol manifested severe signs of crypt loss, suggesting to be the cause of death as can be observed in severe cases of inflammatory bowel disease (Eichele & Kharbanda, 2017). These observations inspired us to test the impact of decreased numbers of colonic PC in inflammation. To this aim, ColVIcre*-Kif3a*^*flx/flx*^ and control mice were treated for one week with DSS (Figure 4A), a commonly used animal model for acute colitis (Wirtz *et al*, 2007)(Hao *et al*, 2015). In fact, ColVIcre-*Kif3a*^*flx/flx*^ mice displayed a significantly higher weight loss than Kif3a^*flx/flx*^ mice at days 7, 8 and 9 after the start of the protocol (Figure 4B). Concurring with the more pronounced weight loss, colons of ColVIcre*-Kif3a*^*flx/flx*^ mice displayed larger areas of crypt loss and architectural irregularities (Figures 4C,D,E). Notably, 4 out of 7 ColVIcre-*Kif3a*^*flx/flx*^ mice, but none of the control animals, exhibited signs of dysplasia (Figure 4F). Moreover, ColVIcre*-Kif3a*^*flx/flx*^ mice displayed an increased presence of F4/80^positive^ macrophages in areas with crypt loss (Figure 4G), whereas numbers of Gr-1^positive^ granulocytes, B220^positive^ B cells and CD3^positive^ T cells were unaltered (supplementary Figure 4). Consistent with the increased susceptibility of ColVIcre*-Kif3a*^*flx/flx*^ mice to DSS-induced colitis we detected decreased numbers of PC on colonic fibroblasts during acute colitis compared to those of untreated mice further underpinning a link between the presence of PC and intestinal inflammation (Figure 4G). Strikingly, we also detected decreased numbers of PC in regions of inflamed tissue from patients with ulcerative colitis, when compared to neighboring normal tissue (Figure 4I).

**Figure 4.**
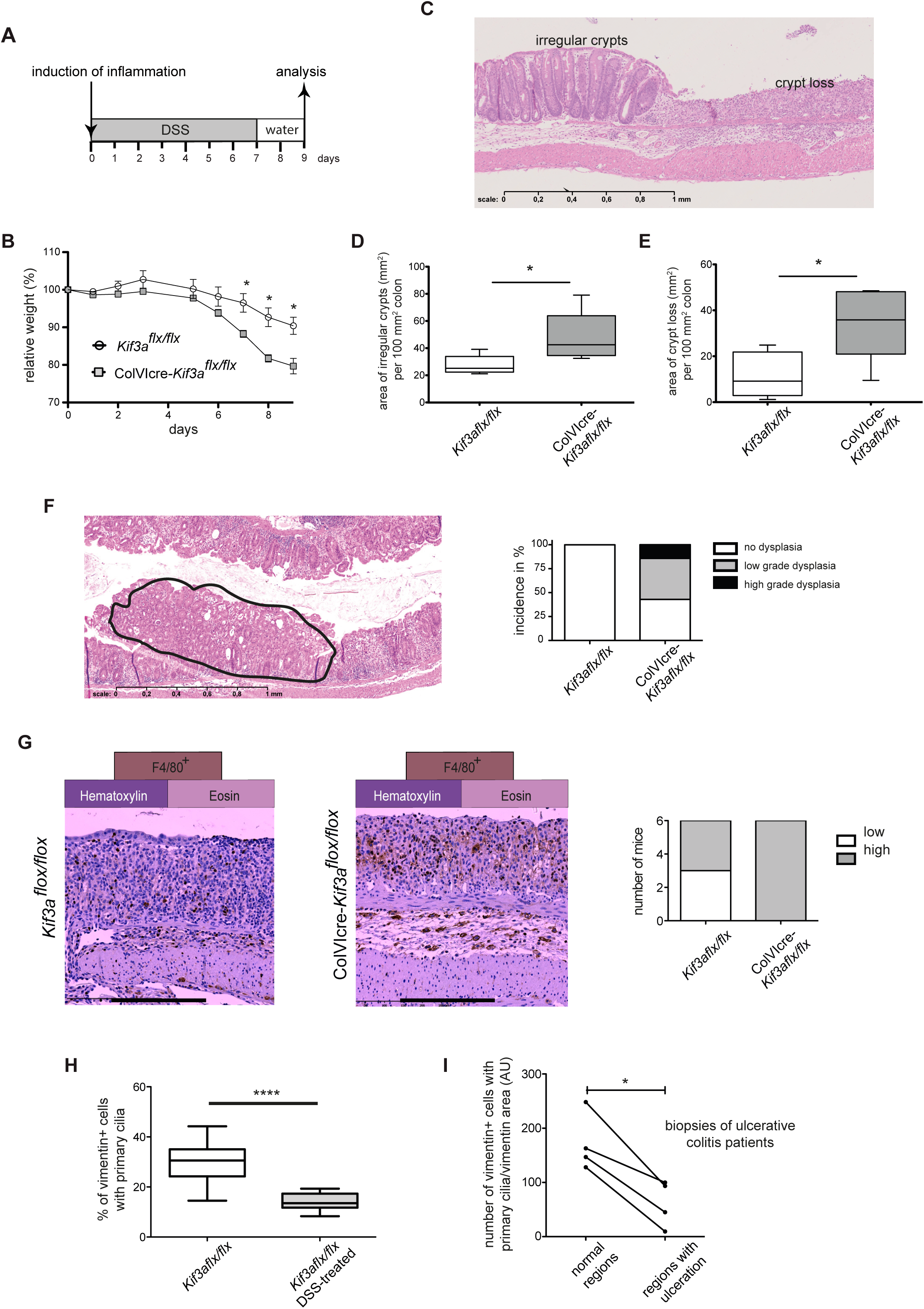
ColVIcre-*Kif3a*^*flx/flx*^ mice are more susceptible to DSS-induced colitis. (A) Experimental timeline for DSS-induced colitis in control (*n* = 6) and ColVIcre-*Kif3a*^*flx/flx*^ (*n* = 6) male mice. (B) Weight development in DSS-treated mice. *p<0.05 by two-tailed unpaired t-tests. (C,D,E) At the end of the protocol described in (A) mice were sacrificed and colons were analyzed by histology for colitis severity including areas of crypt loss (C,D) and architectural irregularities (C,E). Data are presented as Box-and-whisker plots. *p<0.05 by two-tailed unpaired t-test. (F) Incidence of dysplasia in DSS-treated ColVIcre-*Kif3a*^*flx/flx*^ mice. A representative image of a region with dysplasia is shown (circled area with a region of high grade dysplasia on the left, and low grade dysplasia on the right side), as well as the incidence of low and high grade dysplasia in ColVIcre-*Kif3a*^*flx/flx*^ and control mice. (G) Elevated numbers of F4/80^+^ macrophages in colons of DSS-treated *ColVIcre-Kif3a*^*flx/flx*^ mice. Representative images for F4/80 staining in areas with crypt loss are shown. At least 5 fields in the regions of crypt loss were analyzed from each colon of control (n=6) and ColVIcre-*Kif3a*^*flx/flx*^ (n=6) mice. Mean cell numbers were scored as low (<500 cells/mm^2^) or high (>500 cells/mm^2^). Scale bar represent 250µm. (H) Comparison of primary cilia numbers on vimentin^positive^ cells in colons of untreated and DSS-treated wild type mice. DSS-treatment was performed as illustrated in Figure 4A. Primary cilia in DSS-treated mice were depicted in areas of regeneration (14 fields of in average 0.250mm^2^). Significance values were determined by two-tailed unpaired t-tests (****= p<10^−4^). (I) Decreased number of PC in inflamed tissue of patients with ulcerative colitis. Shown are the results of matched-pair analysis for PC expression per vimentin^+^ cells as arbitrary unit (AU, see material and methods). The lines link values obtained for regions of ulceration in the ileum with those of normal regions from the same patient, demonstrating the difference in each pair. *p<0.05 by paired t-test

### ColVIcre-*IFT88*^*flx/flx*^ mice display less colonic PC and increased susceptibility to DSS-induced colitis

Our analysis of ColVIcre*-Kif3a*^*flx/flx*^ mice revealed a link between PC loss on colonic fibroblasts and susceptibility to DSS-induced colitis as well as AOM/DSS-induced carcinogenesis. To validate these observations, we generated a second mouse model for PC loss in colonic fibroblasts by crossing a conditional knockout allele of the intra-flagellar transport protein 88 (*Ift88*^*flx/flx*^) with ColVI-cre mice. Similar to Kif3A, Ift88 is essential for cilia assembly and absence of this protein leads to the loss of PC (Haycraft *et al*, 2007). As expected, ColVIcre-mediated deletion of *IFT88* was only detectable in colonic fibroblasts but not epithelial cells (Supplementary Figure S5), resulting in decreased numbers of PC in vimentin^+^ colonic cells (Figures 5A). ColVIcre-*Ift88*^*flx/flx*^ mice were more susceptible to DSS-induced colitis compared to control animals, as manifested by elevated weight loss, decreased colon size and increased crypt loss (Figures 5B-D). Moreover, male ColVIcre-*Ift88*^*flx/flx*^ mice showed, similar to ColVIcre*-Kif3a*^*flx/flx*^ animals, a significant lower survival rate when exposed to the AOM/DSS colon carcinogenesis protocol (Figure 5E).

**Figure 5.**
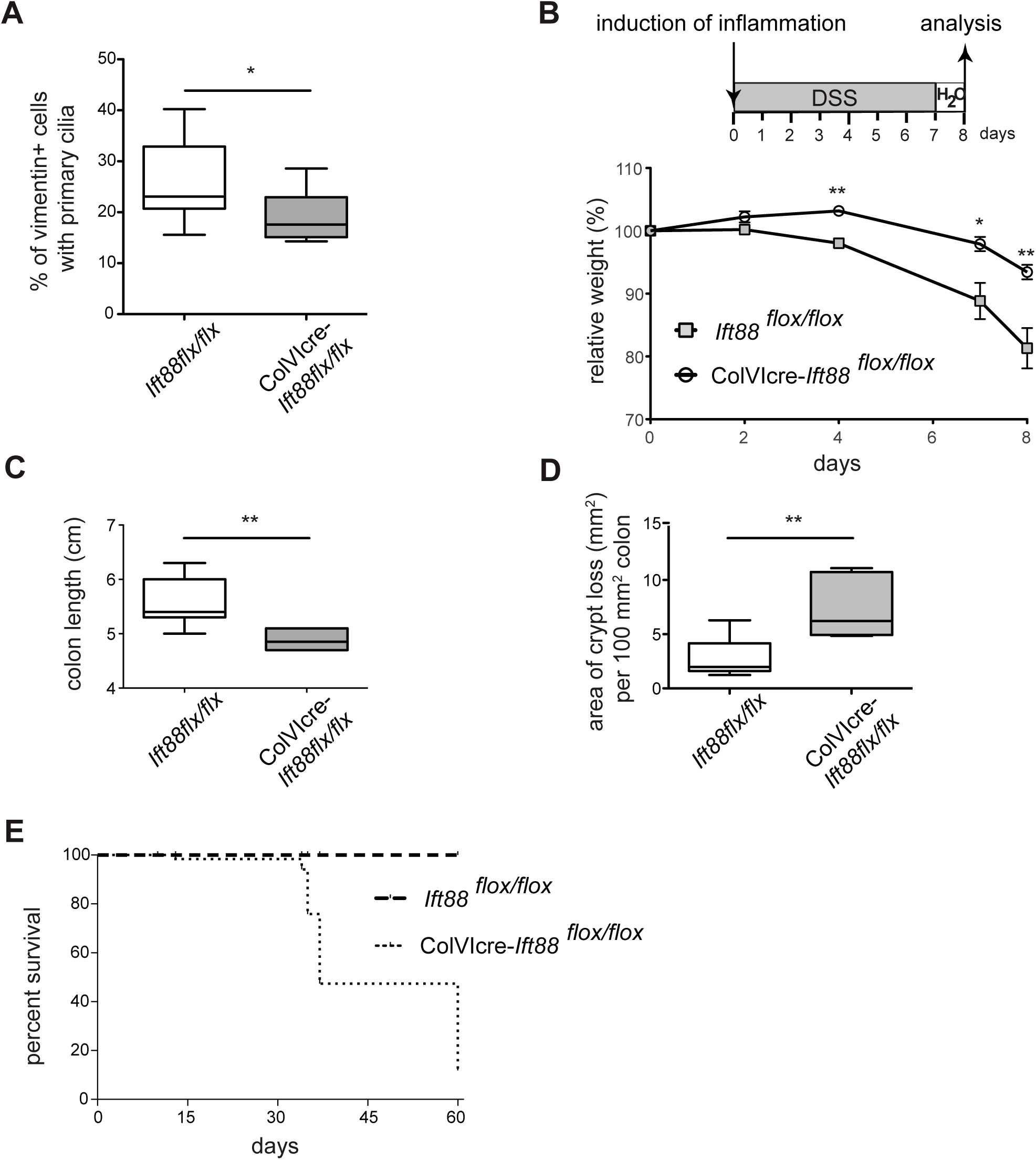
Decreased numbers of colonic primary cilia in ColVIcre-*Ift88*^*flx/flx*^ mice increases the susceptibility to DSS-induced colitis and CAC. (A) Quantitative analysis of PC expression on vimentin^+^ cells in normal colon of *Ift88*^*flx/flx*^ (n=7) *and* ColVIcre-*IFT88*^*flx/flx*^ (n=6) mice. *p<0.05 by two-tailed unpaired t-test. (B) Weight development in DSS-treated ColVIcre-*Ift88*^*flx/flx*^ (n=6) and control (n=7) mice following the indicated timeline. (C,D) At the end of the protocol described in (B) mice were sacrificed and colons were analyzed for length (C), and crypt loss (D). Data are presented as Box-and-whisker plots. *p<0,05 by two-tailed unpaired t-test. (E) Kaplan-Meier survival curves of AOM/DSS-treated male *Ift88*^*flx/flx*^ (n=11) *and* ColVIcre-*Ift88*^*flx/flx*^ (n=12) mice. Colon carcinogenesis was induced as described in Figure 2A.

### Colons from *DSS*-treated PC-deficient mice exhibit an altered regenerative response

We next looked into the molecular alterations associated with the elevated inflammatory response of the PC-deficient mouse strains. A critical role for the prototypic pro-inflammatory cytokine IL-6 in DSS-induced colitis is well established and indeed, we detected elevated IL-6 transcript levels in distal colons from DSS-treated ColVIcre*-Kif3a*^*flx/flx*^ mice as compared to controls (Supplementary Figure S6A). Macrophages can be important producers of IL-6 (Naito *et al*, 2004) and a higher number of F4/80^positive^ macrophages in areas with crypt loss from DSS-treated ColVIcre*-Kif3a*^*flx/flx*^ mice showed detectable Il-6 expression by IHC (Supplementary Figure S6B). In addition to macrophages, fibroblasts and epithelial cells can be significant producers of IL-6 (Waldner *et al*, 2012), and in DSS-treated ColVIcre-*IFT88*^*flx/flx*^ mice both displayed an elevated production of IL-6 as determined by immunofluorescent analysis of colons (Figure 6A). Targets of IL-6 are the transcription factors NFκB and STAT3, which can interact with each other and play an important role in linking inflammation and cancer (Karin & Clevers, 2016)(Taniguchi & Karin, 2018). A more frequent nuclear localization and thus activation of NFκB was detectable in colonic fibroblasts and epithelial cells of DSS-treated ColVIcre-*IFT88*^*flx/flx*^ mice, whereas higher levels of nuclear phospho-STAT3 were only evident in vimentin^+^ fibroblasts (Figures 6B,C). Strikingly, DSS-treated colons of ColVIcre-*IFT88*^*flx/flx*^ animals showed significantly lower expression levels of the transcription factor HES1 (Figure 6D), an important downstream target of Notch signaling, a key pathway for the regeneration of intestinal epithelial cells during DSS-induced colitis (Okamoto *et al*, 2009). This finding concurs with the decreased transcript levels of Hes1 found in the distal region of colons from colitic ColVIcre-*IFT88*^*flx/flx*^ mice (Supplementary Figure S6A).

**Figure 6.**
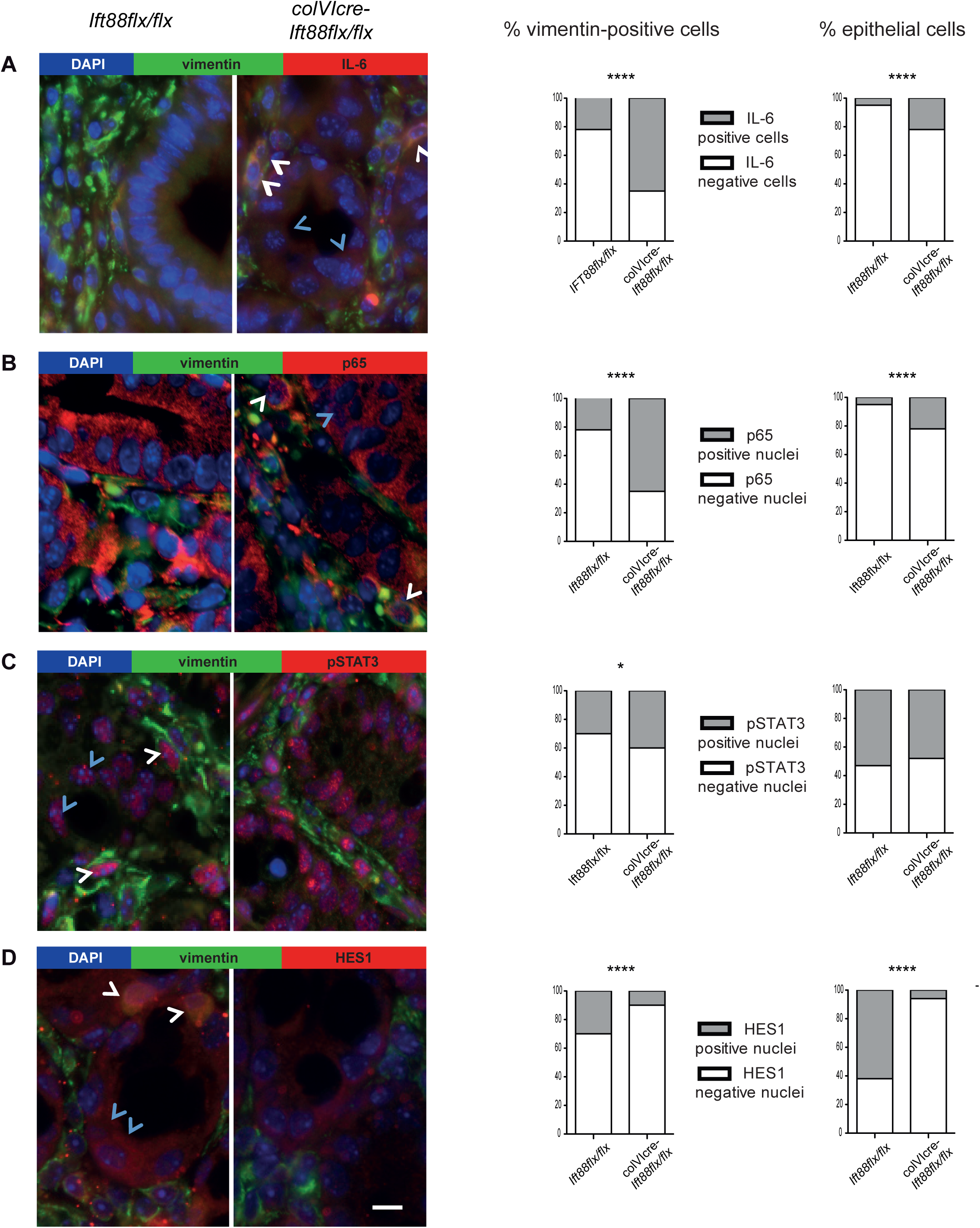
Colons from DSS-treated ColVIcre-*IFT88*^*flx/flx*^ display an altered regenerative response. (A, B, C, D) Representative images and corresponding quantification of vimentin^positive^ fibroblasts (in green) and vimentin^negative^epithelial cells (identified by histological criteria) labeled for IL-6 (A), p65 (B), pSTAT3 (C) and HES1 (D) (all in red) in colons of *Ift88*^*flx/flx*^ and ColVIcre-*Ift88*^*flx/flx*^ mice treated with DSS as described in Figure 4A. White arrow heads indicate IL-6 (A), nuclear staining for p65 (B), pSTAT3 (C) and HES1 (D) for fibroblasts, blue arrow heads the respective labeling for epithelial cells. Nuclei were stained with DAPI (in blue). Scale bars represent 100 µm. *p<0.05; ****p<10^−4^ was calculated by chi-squared test on absolute numbers and is presented as % of cells positive for the indicated marker within vimentin^positive^ and epithelial cells, respectively.

### Decreased numbers of PC in human CRC biopsies

To investigate whether PC are also present in human colons we performed IHC analysis of three biopsies of healthy donors. Indeed, PC were detectable mostly on vimentin^+^ fibroblasts in the lamina propria and only on few epithelial cells (Figure 7A, Supplementary Figures S7A,B). We next analyzed the presence of PC on tumor tissues of 28 CRC patients at different stages of disease. Sections of tumor tissues of all four stages harboured decreased numbers of PC on vimentin^positive^ colonic fibroblasts when compared to the respective peri-tumoral region as well as normal colon (Figure 7B). Of note, the tissue sections of the stage 4 samples did not contain adjacent normal tissue and were thus compared with the mean calculated from the analysis of peritumoral tissue of stage 1-3 patients. Moreover, we noted that vimentin^positive^ regions were enlarged in the tumoral as compared to the peritumoral regions of several samples. Hence, we analyzed PC expression in regions containing exclusively vimentin^positive^ cells to convey a more unbiased approach. Notably, the latter analysis confirmed the decreased number of PC in tumor as compared to peritumoral areas (Figure 7C).

**Figure 7.**
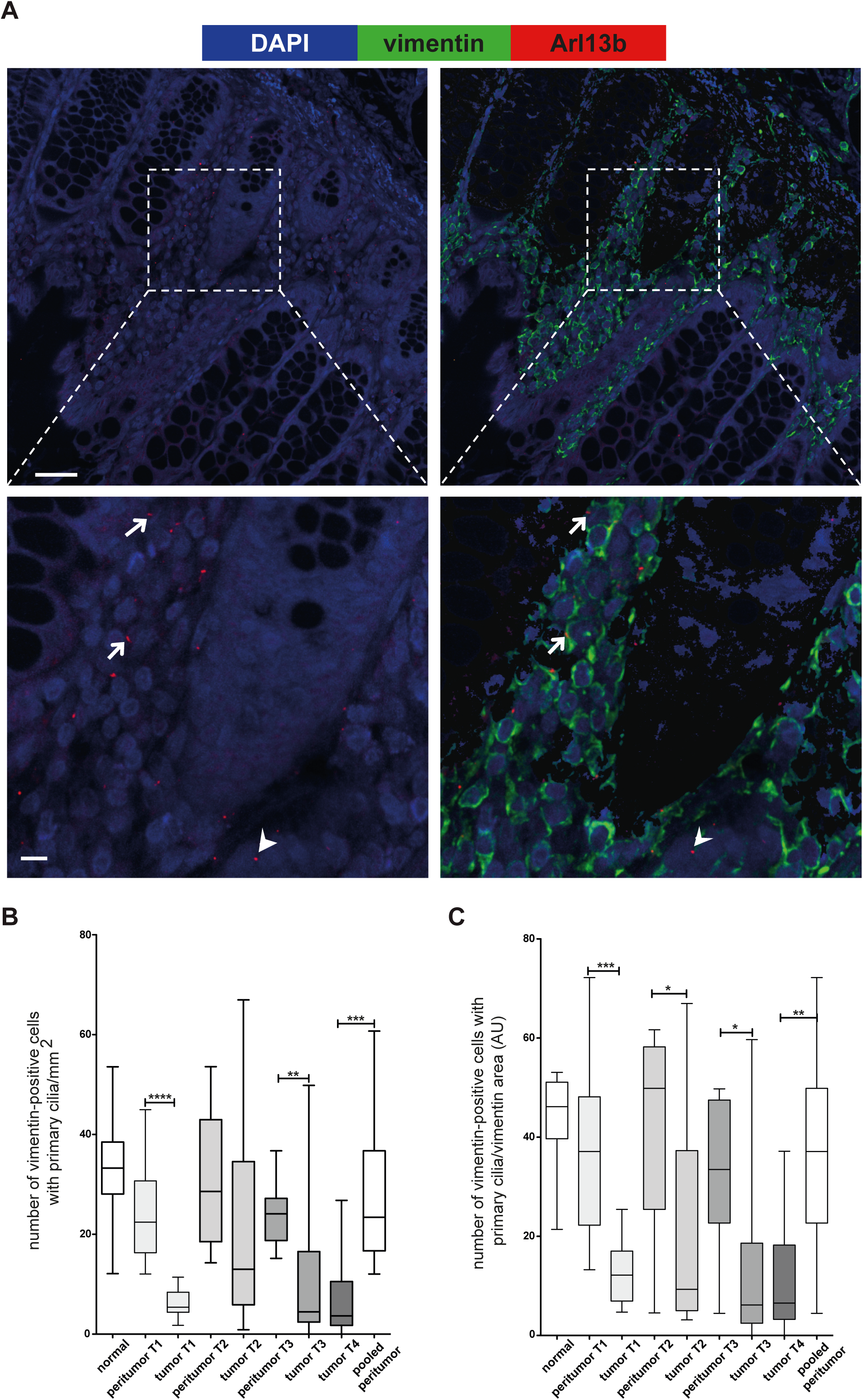
Decreased number of primary cilia in tumoral tissues of CRC patients compared to peritumoral regions. (A) Representative image of vimentin^+^ fibroblasts (in green) expressing PC (identified using Arl13b, in red) in a normal colon. Nuclei were stained with DAPI (in blue). Most PC are detected on vimentin^+^ cells (arrow), and few on vimentin-negative epithelial cells (arrow head). Scale bars represent 50 µm in the upper and 10 µm in the lower panel. (B,C) Quantitative analysis of PC presence on tumoral and peri-tumoral regions of colons from CRC patients (n=28) at tumor (T) stages 1 to 4. The panels represent quantification of PC expression (B) of vimentin^positive^ cells per mm^2^ and (C) per vimentin^positive^ cells as arbitrary unit (AU, see material and methods). Data are presented as Box-and-whisker plots. *p<0,05, **p<0,01, ****p<10^−4^ by two-tailed unpaired t-test. Images were acquired on an Andor Dragonfly Spinning Disk Confocal microscope and analyzed with the Imaris software. Quantification of PC and Vimentin^+^ fibroblasts was done on at least 5 random fields (about 1mm^2^ each) per sample using ImageJ software.

## Discussion

Various reports associate PC with tumorigenesis, but their role in tumor development appears to vary between different types of tumors. Reduced numbers of PC have been described for breast and pancreatic cancer, as well as melanoma (reviewed in (Liu *et al*, 2018)(Eguether & Hahne, 2018)), whereas PC were reported to be maintained in about half of the tested biopsies of medulloblastoma and basal cell carcinoma (BCC) patients (Han *et al*, 2009)(Wong *et al*, 2009). This dichotomy is underlined by the observation in mouse models that, depending on the nature of the initiating oncogenic event, cilia ablation facilitates or blocks medulloblastoma as well as BCC tumor formation (Han *et al*, 2009)(Wong *et al*, 2009).

Here, we report that numbers of PC decrease during colon carcinogenesis in mice. Intriguingly, this finding in mice concurs with the lower number of PC in tumor areas of stage 1-4 CRC patients, when compared to peritumoral regions as well as healthy controls. We initiated a multicenter study to confirm this finding on a larger cohort of CRC samples and to evaluate whether CMS subtype-specific PC patterns exist. In line, a recent report described a correlation between the frequency of colonic PC and disease outcome in CRC patients (Dvorak *et al*, 2016). The authors found a significantly longer overall survival of CRC patients with a higher frequency of PC in the colon, concurring with our observations in mice. Nevertheless, this study did not assess which cells in the colon do express PC.

We found that in murine as well as human colon, mostly vimentin^positive^ stromal cells carry PC, which are, however, detectable only on a part of those cells. To investigate consequences of PC loss in colonic fibroblasts, we crossed the two commonly used mouse strains to study PC, i.e. *Kif3a*^*flx/flx*^ and *Ift88*^*flx/flx*^, to ColVIcre-transgenic mice, targeting mesenchymal cells including subsets of colonic fibroblasts (Armaka *et al*, 2008)(Koliaraki *et al*, 2015). Accordingly, the number of PC was significantly lower in colons of both ColVIcre-*Kif3a*^*flx/flx*^ and ColVIcre-*Ift88*^*flx/flx*^ mice compared to control animals, in agreement with the requirement of KIF3a and IFT88 for maintenance of PC. The decreased number of PC in the colon of these mice did not perturb the colonic architecture. This corroborates our previous observation in *Ttll3*-deficient mice, in which the number of PC is similarly diminished without affecting colon architecture and homeostasis (Rocha *et al*, 2014).

It is well established that colonic fibroblasts cooperate with epithelial cells during colon carcinogenesis (Vermeulen *et al*, 2010). We studied the consequences of the loss of PC on colonic fibroblasts by exposing ColVIcre-*Kif3a*^*flx/flx*^ as well as ColVIcre-*IFT88*^*flx/flx*^ animals to chemically induced colitis-associated colon carcinogenesis (AOM/DSS model). Both mutant mouse strains displayed increased dysplasia suggesting that the partial loss of PC on colonic fibroblasts promotes tumorigenesis. In addition to their role in PC formation, KIF3a and IFT88 may promote cilia-independent functions, such as spindle orientation or mother centriole appendage formation (Kodani *et al*, 2013)(Delaval *et al*, 2011). Therefore, we decided to study both mutant mice deficient of PC to allow a conclusion whether a potential phenotype is truly cilia-dependent. The complementary results obtained in ColVIcre-*Kif3a*^*flx/flx*^ and ColVIcre-*Ift88*^*flx/flx*^ animals allow to conclude that decreased PC numbers are at their origin.

Interestingly, two recent articles described different roles for subsets of colonic fibroblast during CAC in mice (Pallangyo *et al*, 2015)(Koliaraki *et al*, 2015). In both studies, the role of IKKβ in CAC was investigated upon tissue-specific deletion in subsets of stromal cells in the colon using either colagenase1a2 (*Col1a2*)- or *ColVI*-Cre driven DNA recombination. In the former model tumor promotion was observed, while the latter resulted in decreased tumor formation. A likely explanation for these findings might be that the *col1a2*- and *colVI*-promoters target different stromal cell subsets, which may have distinct functions (Fordham & Sansom, 2015). In fact, it appears that intestinal stromal cells consist of several subtypes that are not well defined yet (Roulis & Flavell, 2016). In addition, it has been suggested that resting fibroblasts are capable to differentiate into distinct subsets that execute different functions (Kalluri, 2016). This is well illustrated by a recent report describing the presence of two different subsets, i.e. myofibroblasts and inflammatory fibroblasts, in pancreatic ductal adenocarcinoma (Öhlund *et al*, 2017).

We found that deficiency of PC on colonic fibroblasts renders ColVIcre-*Kif3a*^*flx/flx*^ as well ColVIcre-*Ift88*^*flx/flx*^ mice more susceptible to DSS-induced colitis. Accordingly, decreased numbers of PC were detected in the colons of colitic mice. Moreover, the decreased number of PC detected in inflamed tissue areas of patients with ulcerative colitis corresponds with our observations in mice. Analysis of the distal parts of colons isolated from DSS-treated animals revealed elevated IL-6 transcript levels in ColVIcre-*Kif3a*^*flx/flx*^ mice, which correlates with elevated numbers of macrophages that are well-established producers of IL-6 during inflammation in areas of crypt loss in mutant mice. Notably, in areas without crypt loss increased numbers of IL-6 expressing colonic fibroblasts were detectable in ColVIcre-*Kif3a*^*flx/flx*^ mice concurring with reports that colonic fibroblasts have the capacity to produce significant levels of IL-6 (Lin *et al*, 2013)(Koliaraki *et al*, 2015)(Shalapour & Karin, 2015). The increased production of pro-inflammatory cytokines was linked to activation of NF-κb and Stat3 signaling in PC-deficient colonic fibroblasts derived from ColVIcre-*Kif3a*^*flx/flx*^ mice, in agreement with a report that reduced NF-κb signaling in ColVIcre targeted colonic fibroblasts dampens IL-6 production and thus inflammation in DSS-induced colitis (Koliaraki *et al*, 2015). Reports associating altered PC numbers with inflammation *in vivo* are still limited. A recent work from Reiter and colleagues describes that removal of PC on endothelial cells is dispensable for vascular development in mice (Dinsmore & Reiter, 2016). Removal of endothelial PC, however, was found to promote atherosclerosis in mice that correlated with an increased expression of inflammatory genes including notably IL-6 and NF-κb (Dinsmore & Reiter, 2016), thus concurring with our observations.

A concept is emerging that inflammation is an important driver in the regeneration of damaged tissue, by inducing hyperproliferation of epithelial tissue triggered by pro-inflammatory cytokines such as IL-6 (Kuhn *et al*, 2014)(Karin & Clevers, 2016). Excessive production of pro-inflammatory cytokines, however, can also be deleterious and promote tumor formation (Todoric *et al*, 2016). In fact, we observed an increased incidence of dysplasia in DSS-treated ColVIcre-*Kif3a*^*flx/flx*^ mice. DSS is not genotoxic, but intestinal inflammation was shown to induce DNA damage and thus promoting dysplasia (Westbrook *et al*, 2009). Therefore, the augmented inflammatory response in DSS-treated ColVIcre-*Kif3a*^*flx/flx*^ mice is most likely related to the observed incidence of dysplasia.

Several signaling pathways regulate the wound-healing process including NF-kB, and STAT3 (Karin & Clevers, 2016). In addition, intact Notch signaling is required for the regenerative response of intestinal epithelia upon DSS-treatment (Okamoto *et al*, 2009)(Fazio & Ricciardiello, 2016). Colonic epithelial cells in DSS-treated ColVIcre-*Kif3a*^flx/flx^ mice displayed significantly lower expression levels of the Notch target HES1, suggesting that impaired Notch signaling contributes to the disturbed wound healing process detected in these mice. Indeed, absent or very weak expression of HES1 was found in sessile serrated adenoma/polyp (SSA/p), a precursor lesion for colorectal carcinoma (Cui *et al*, 2016). Moreover, recent reports describe an inflammation and tumor-suppressive function of Notch/Hes1 signaling in the colonic epithelium of patients with ulcerative colitis or CRC, respectively (Wang *et al*, 2017)(Tsuchiya *et al*, 2017; Prossomariti *et al*, 2017).

Taken together, decreased numbers of PC on colonic fibroblasts in mice has no consequence in steady-state but enhances the inflammatory response in acute colitis and promotes chemically induced colon carcinogenesis. The observations made in mice concur with the corresponding human pathologies, i.e. colitis and CRC, as those display decreased numbers of PC on intestinal fibroblasts.

## Material and Methods

### Animal experimentation

Mouse experiments were performed in strict accordance with the guidelines of the European Community (86/609/EEC) and the French National Committee (87/848) for care and use of laboratory animals. To study PC we used *Kif3a*^*flx/flx*^ and *Ift88*^*flx/flx*^ mice which have been previously described (Marszalek *et al*, 1999)(Haycraft *et al*, 2007). Tissue specific knock-out mice were obtained by crossing with ColVIcre-transgenic strain. The latter was obtained by G. Kolias (Armaka *et al*, 2008)(Koliaraki *et al*, 2015). Mice were maintained on C57BL/6 genetic background, and experimental groups contained littermates that were caged together according to gender. Genotyping was done as described before (Marszalek *et al*, 1999)(Haycraft *et al*, 2007)(el Marjou *et al*, 2004)(Armaka *et al*, 2008). ColVIcre-*Kif3a*^*flx/flx*^ and ColVIcre-*Ift88*^*flx/fl*^ mice were fertile, born at expected mendelian ratio and displayed no overt intestinal phenotype. Mice used for the experiments described were used at age of 8-12 weeks.

### Patient samples

Formalin-fixed paraffin-embedded CRC samples of different grades were obtained from the institutional biobank of the University Hospital of Liege, Belgium following the approval of the Institutional Ethics Committee No. 2009/69. According to Belgian law, informed consent was not necessary because all patients are informed that their residual surgical material can be used for research unless they opt-out. Details of CRC samples obtained from the University Hospital (CHU) of Liège are listed in Appendix Table S1. The cohort of patients with ulcerative colitis from the Hospital Saint Antoine (Paris) was declared under the number #CNIL1104603 and all patients (two men, 38 and 43 years old, and two women, 25 and 57 years old) gave consent to this research.

### Histology and immunohistochemistry

Organs were fixed in formalin solution for 24h. Histological examination was performed on paraffin-embedded sections stained with hematoxylin and eosin. Immunohistochemistry was performed on formalin-fixed and paraffin-embedded tissues cut into 4-µm sections. After blocking of non-specific binding (with TBS-10%goat serum-5%BSA-5%milk-0,3%triton), samples were incubated with primary antibody (see Table 1) for 1 h at room temperature or overnight at 4°C and avidin/biotin or polymer horse radish peroxidase kits for primary mouse or rabbit antibodies (Vector Laboratories) were used for detection. For immunofluorescence analysis, samples were incubated with primary antibodies for 1h and revealed with fluorescent-labeled secondary antibodies (Vector Laboratories). DNA was stained with 20 µg/ml 4’,6’-diamidino-2-phenylindole (DAPI). For the detection of PC we followed a previously described protocol that allows detection of PC on paraffin embedded tissue (Hassounah *et al*, 2013).

**Table 1:**
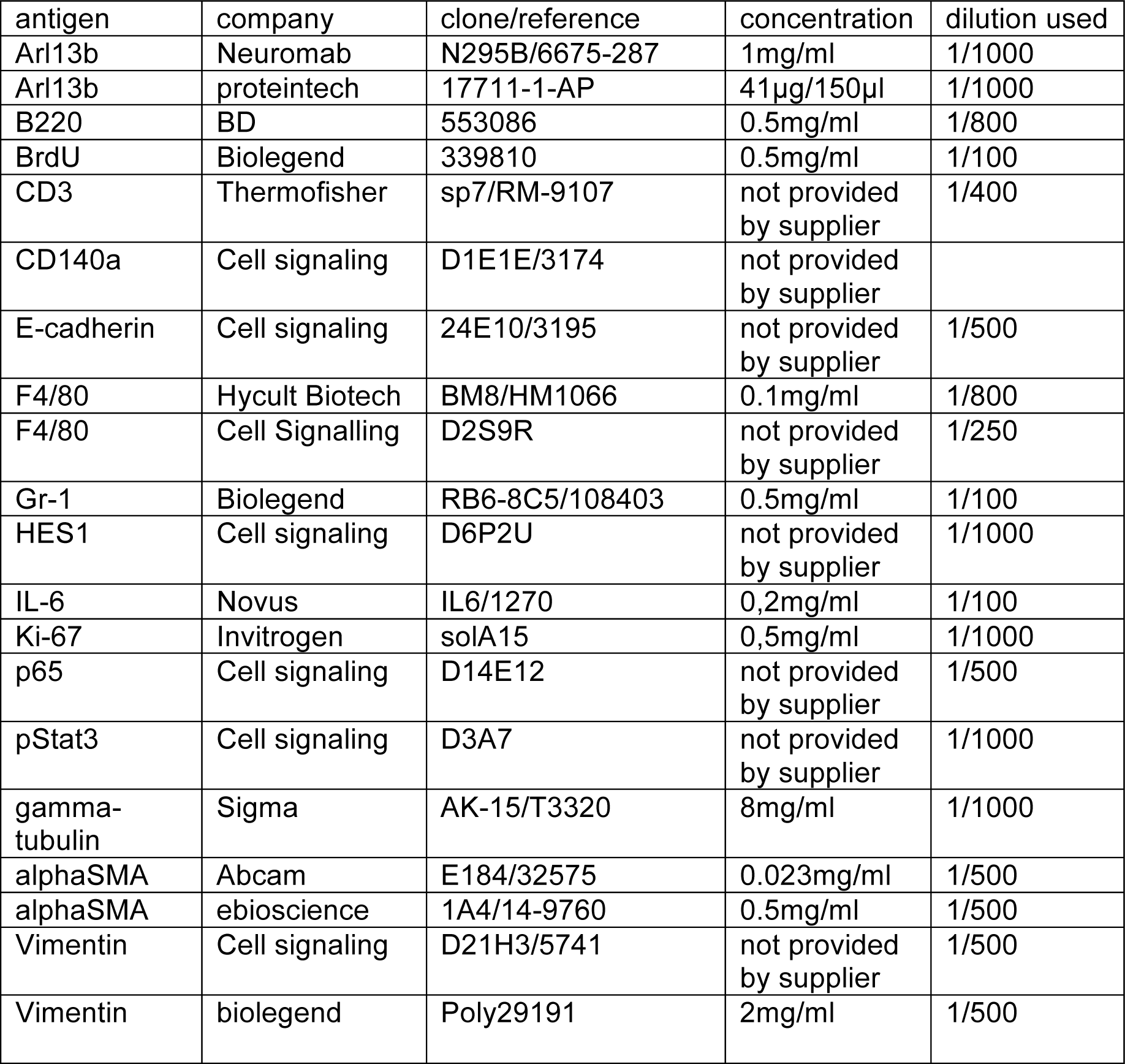
Antibodies used in IHC on paraffin embedded tissues.

### DSS-induced colitis and colitis associated carcinogenesis (CAC)

To induce colitis, 8-12 week-old mice were treated for 7 days with 2.5% (w/v) dextran sodium sulfate (DSS; TdB Sweden) added in the drinking water. Mice were sacrificed at indicated time points. The weight of the mice was daily followed and in case weight loss was more important than 20% mice were sacrificed.

For colitis-associated carcinogenesis, mice were intraperitoneally injected with azoxymethane (AOM), followed by three cycles of 2.5% (w/v) DSS administered in the drinking water (Figure 2A). In between the cycles the mice received no treatment for 2 weeks. For histological analysis the entire colon was prepared according to the Swiss roll procedure, fixed with formaldehyde and embedded in paraffin. Four µm sections were stained with hematoxylin and eosin, Alcian blue or deparaffinized and subsequently incubated with the primary antibody described below. Histological grading of AOM/DSS-induced tumors was determined with blinded genotype according to the described classification of intestinal neoplasia (Washington *et al*, 2013).

### BrdU incorporation proliferation assay

12-week-old mice were intraperitoneally injected with 100 µg Bromodeoxyuridine (BrdU) per gram body weight. Colons were dissected 2 h or 5 days following injection. Proliferating cells were detected with anti-BrdU antibody (Biolegend), and BrdU^+^ cells were quantified from at least three animals per condition.

### Cell culture

Protocols for isolation of intestinal epithelial cells (IEC) and fibroblasts (InF) were modified according to Roulis *et al.* (Roulis *et al*, 2014). Briefly, colons were isolated from mice, flushed with PBS/2% FCS, and then opened longitudinally and processed as follows. Intestinal pieces were cut into small pieces and incubated in IEC buffer (PBS/2% FCS/1 mM EDTA/1 mM DTT) at 37°C for 45 minutes under continuous shaking to releases IEC. For the isolation of InF the colon was treated as described to separate IEC, washed, and further processed by treatment with 3 mg/ml Collagenase type I (Sigma) and 0.1 mg/ml Dispase (Dominique Dutscher SA, France) in DMEM for 30 min at 37°C. Cells were filtered through a 70-µm strainer, washed with PBS/2% FCS and subsequently analyzed.

### Microscopy and imaging

Histological slides were scanned using Nanozoomer 2.0 HT scanner with a 40x objective, and visualized with NDP.view2 viewing software (Hamamatsu). Fluorescent images were acquired on a brightfield microscope (Leica) using Metamorph software, inverted Confocal SP5 (Leica) using the Leica LAS AF software or an Andor Dragonfly Spinning Disk Confocal microscope employing Imaris software. Quantification of PC and Vimentin^positive^ fibroblasts was done on at least 5 random fields (about 1mm^2^ each) per sample using Imaris and ImageJ softwares. Images were processed with ImageJ. Images were assembled and adjusted with Adobe Photoshop/ Illustrator. Counting of PC was performed using z-stack acquisition. Co-staining with cell markers was confirmed on distinct layers. In analyses of human biopsies PC numbers were calculated either per mm2 or per Vimentin-stained area using arbitrary unit (AU corresponding to 10000 square pixel) of vimentin^positive^ cells.

### RT-PCR and qRT-PCR

RNA was extracted from homogenized mouse organs or from cells using TRIzol reagent (Euromedex) following the standard protocol. RNA was translated to cDNA with SuperscriptIII reverse transcriptase (Invitrogen) using Random Hexamers (Invitrogen). Quantitative RT-PCR was applied under standard conditions using SYBR Green (Roche) on a LightCycler 480 (Roche). The relative mRNA expression levels of each gene were expressed as the N-fold difference in target gene expression relative to the *TBP* gene.

### Statistical analysis

Statistical analysis was performed employing GraphPad Prism version 5. More precisely, we first validated normal distribution of values (using KS normality test, D’Agostino - Pearson omnibus normality test, Shapiro-Wilk normality test). Then, values of the different groups were analyzed by unpaired t test if the variances of the groups were significantly different and unpaired t test with a Welch’s correction if the variances were significantly different. If values were not normally distributed, a non-parametric Mann-Whitney test was used.

## Acknowledgments

We are grateful to the excellent platforms in Montpellier, i.e. RIO imaging platform, the histology and animal experimentation platforms RHEM and RAM, as well as the IGMM mouse facility. Many thanks to Thierry Gostan and the platform SERANAD for help on the statistical analysis and Prof. BE Clausen for carefully reading the manuscript. This work was supported by INCA (project 2014-215) to MH and CJ, and by the SFNGE (Grant FARE) to MS and TE. Authors are thankful to Dr. Stephanie Gofflot, Biobank CHU Liege for providing clinical information.

## Author’s contribution

RT, CP, RL, TE, HT, JB, CM, SP conducting experiments, acquiring data and analyzing data. AT, MS, PS, VC, PD, CJ, MP, BL, VP, CJ, VP, MH: designing research studies and analyzing data.

MH: writing manuscript.

## Figure legends

**Supplementary Figure 1.** Primary cilia are expressed by few E-cadherin^+^ colonic epithelial cells.

Co-staining was performed for E-cadherin (red) and for primary cilia with Arl13b (arrowhead). Different z-stack slices are shown to validate the co-expression of Arl13b and E-cadherin (slices 13-19 represent 1.2µm thickness). Nuclei were stained with DAPI (in blue). Scale bar represent 10 µm.

**Supplementary Figure 2**

(A) Genotyping of ColVIcre-*Kif3a*^*flx/flx*^ mice displaying *Kif3a*-depletion (ko) in the floxed (flx) alleles in the genomic DNA of total colon and colonic fibroblasts, but not colonic epithelial cells.

(B) Quantitative analysis of primary cilia expression on CD140^+^ cells in the distal, transverse and proximal regions (illustrated by the image on the right hand side) of normal colon of *Kif3a*^*flx/flx*^ and ColVIcre-*Kif3a*^*flx/flx*^ mice as described for Figures 3C and D. ****p<10^−4^ by two-tailed unpaired t-test.

**Supplementary Figure 3**

Relative weight of control (*n* = 7) and ColVIcre-*Kif3a*^*flx/flx*^ (*n* = 7) female mice exposed to the AOM/DSS protocol as described in Figure 2A. *p<0.05, **p < 0.01 by two-tailed unpaired t-test at indicated time points.

**Supplementary Figure 4**

Numbers of Gr1^+^ granulocytes, B220^+^ B cells and CD3^+^ T cells are similar in colons of DSS-treated ColVIcre-*Kif3a*^*flx/flx*^ mice (see Figure 4A). Representative images are shown. At least 5 fields in the regions of crypt loss were analyzed from each colon of control (*n* = 6) and ColVIcre-*Kif3a*^*flx/flx*^ (*n* = 6) mice. Mean cell numbers were scored as low (<500 cells/mm^2^) or high (>500 cells/mm^2^). Scale bar represents 250µm.

**Supplementary Figure 5**

Genotyping of ColVIcre-*Ift88*^*flx/flx*^ mice showing *Ift88*-depletion (ko) in the floxed (flx) alleles in the genomic DNA of colonic fibroblasts, but not colonic epithelial cells.

**Supplementary Figure 6**

Elevated production of IL-6 in colons from DSS-treated ColVIcre-*Ift88*^*flx/flx*^

(A) Transcript levels of Il-6 and Hes1 in DSS treated *Kif3a*^*flx/flx*^ and ColVIcre-*Kif3a*^*flx/flx*^ mice (according to the protocol shown in Figure 5B). Three independent mRNA samples were analyzed by qRT-PCR, and mean values standardized to expression of the house keeping gene *Tbp* are shown. Error bars represent SEM.

(B) Elevated numbers of IL-6-expressing F4/80^+^ macrophages in *ColVIcre-Kif3a*^*flx/flx*^ mice treated with DSS as described in Figure 4. At least 5 fields in the regions of crypt loss were analyzed. Scale bar represent 100 µm. *p<0,05 was calculated by chi-squared test.

**Supplementary Figure 7**

Representative images of the tissue of a stage I CRC patient showing decreased number of primary cilia in tumoral (A) versus peritumoral (B) tissues. PC are indicated by arrows in vimentin-positive fibroblasts, arrow head in epithelial cells and stars in endothelial cells with elongated nucleus. Scale bars represent 50 µm and 10 µm in the upper and lower panel respectively.

**Table S1.** CRC patients Characteristics (n = 28)

| Variable | Value (%) |
| --- | --- |
| Age at diagnosis (yr) |  |
| Median | 66 |
| Range | 42-82 |
| Gender |  |
| Male | 13 (46.4) |
| Female | 15 (53.6) |
| Tumor Location |  |
| Colon | 26 (92.9) |
| ascendant | 15 (53.6) |
| transverse | 2 (7.1) |
| descendent | 2 (7.1) |
| sigmoid | 7 (25) |
| Rectosigmoid | 2 (7.1) |
| Tumor Stage |  |
| I | 6 (21.4) |
| II | 6 (21.4) |
| III | 8 (28.6) |
| IV | 8 (28.6) |
| Metastasis (when data available) |  |
| yes | 6 (21.4) |
| no | 16 (54.1) |

