## supplementary figures for "Partial loss of colonic primary cilia promotes inflammation and carcinogenesis"

### Supplementary figure 1

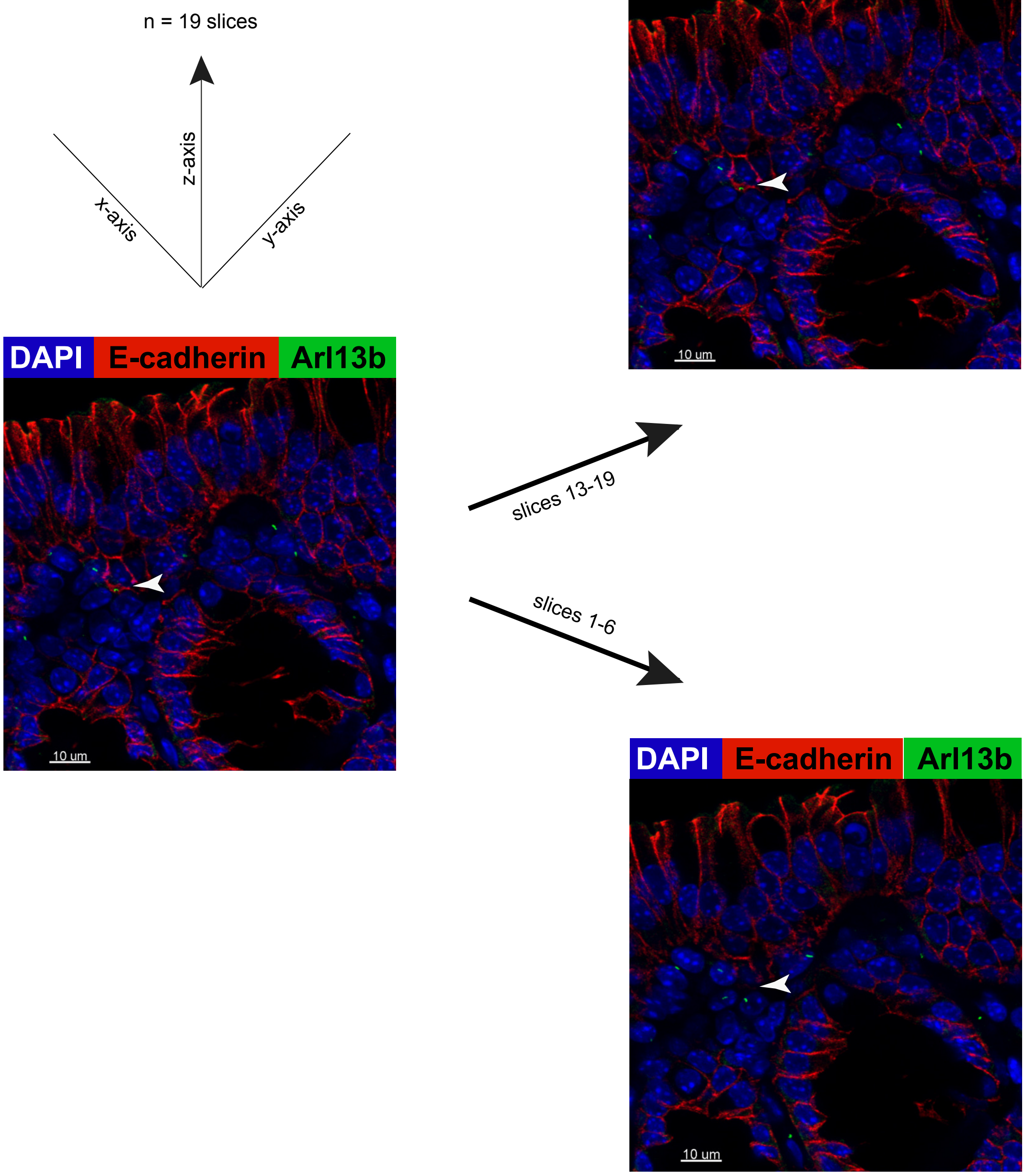

### Supplementary figure 2

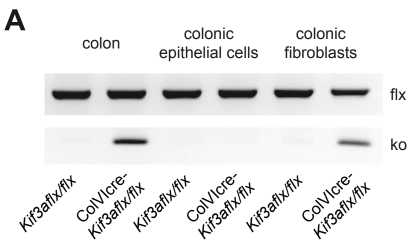


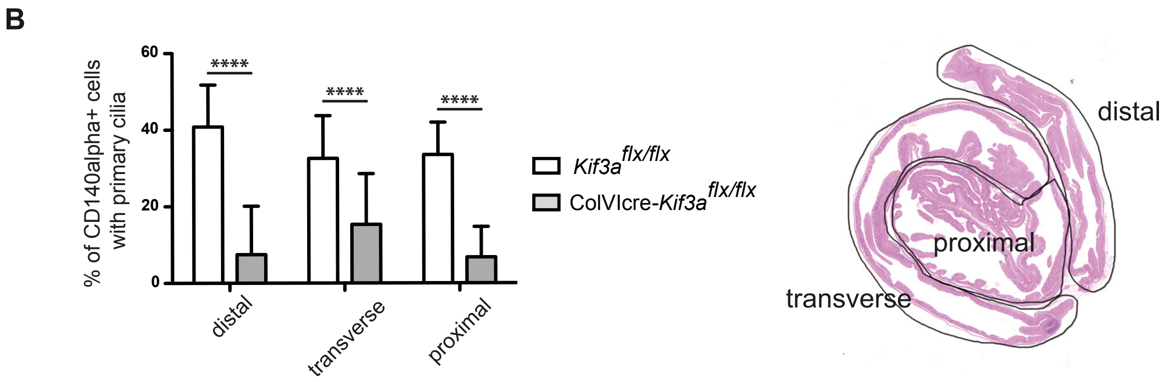

**Supplementary figure 3**

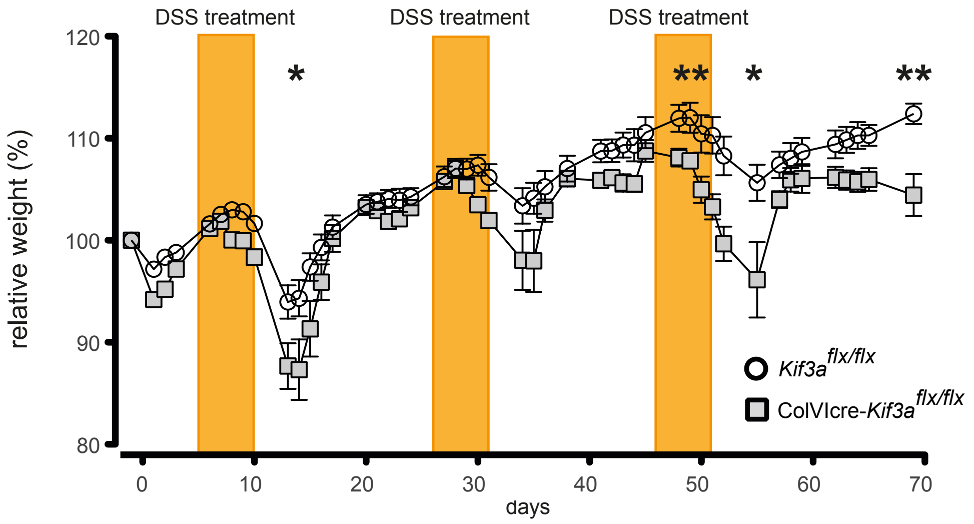

**Supplementary figure 4**

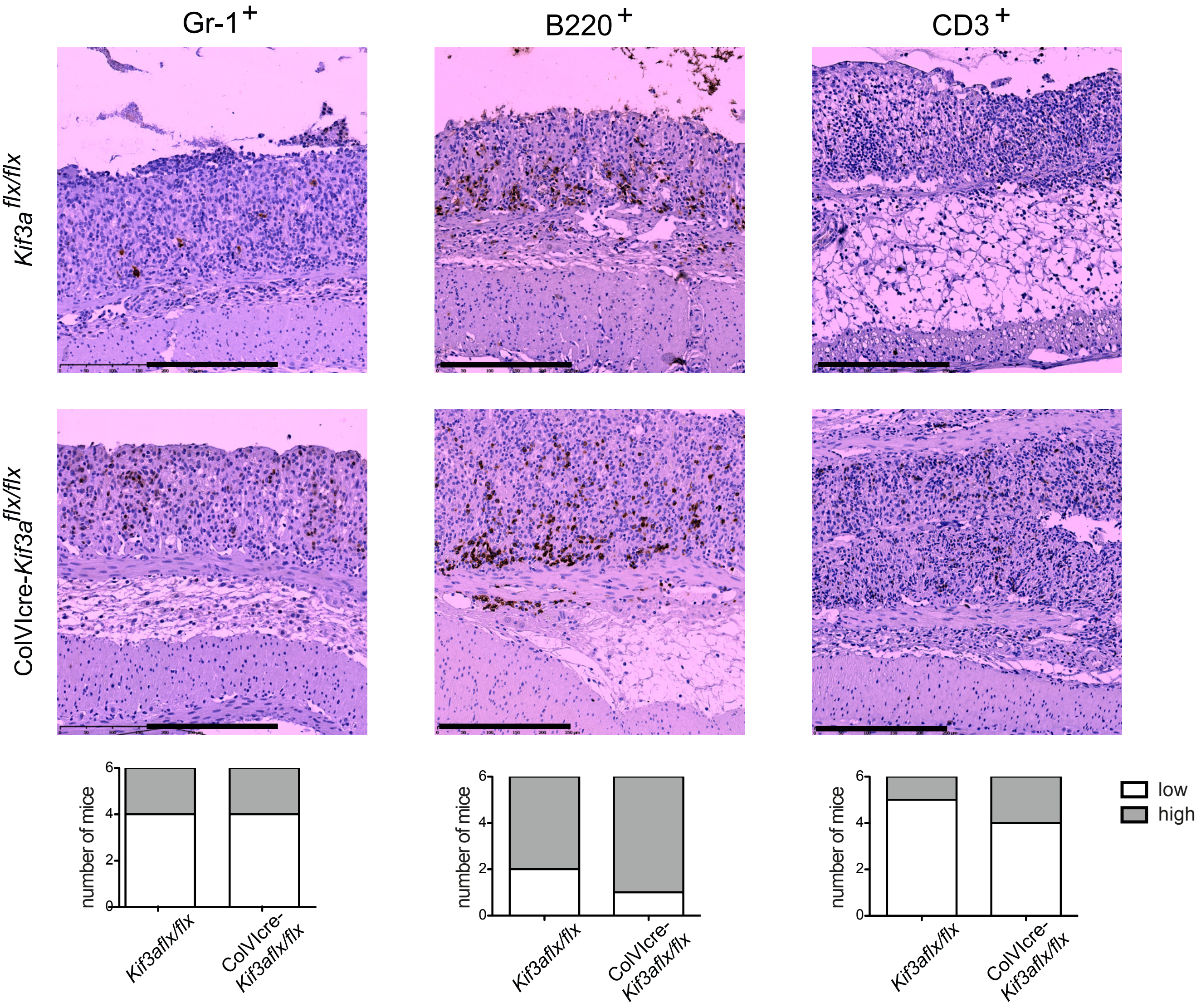

### Supplementary figure 5

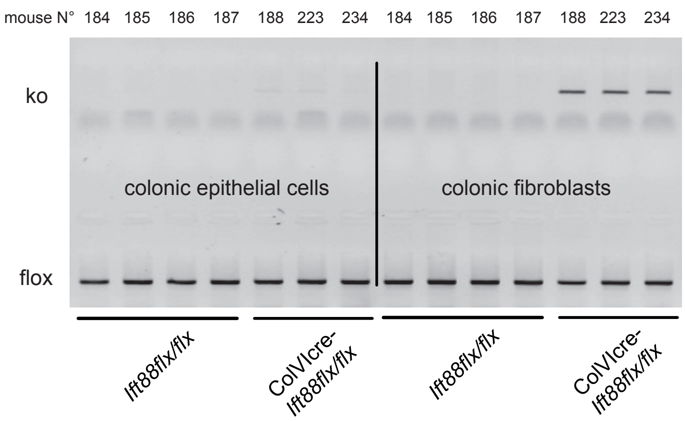

### Supplementary figure 6

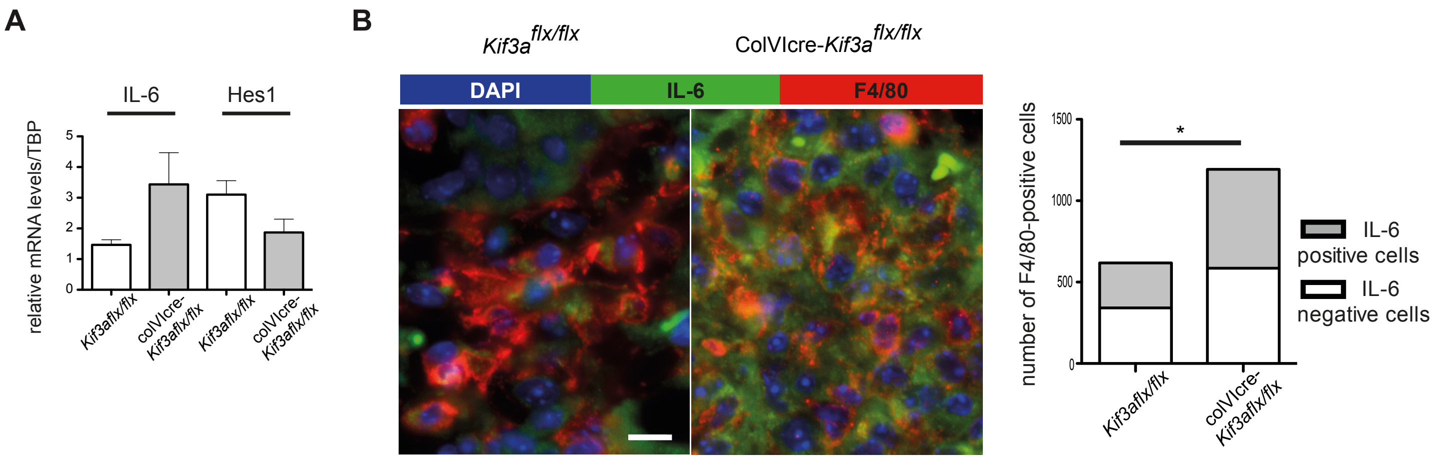

**Supplementary figure 7A**

**Tumoral tissue (Stage 1)**

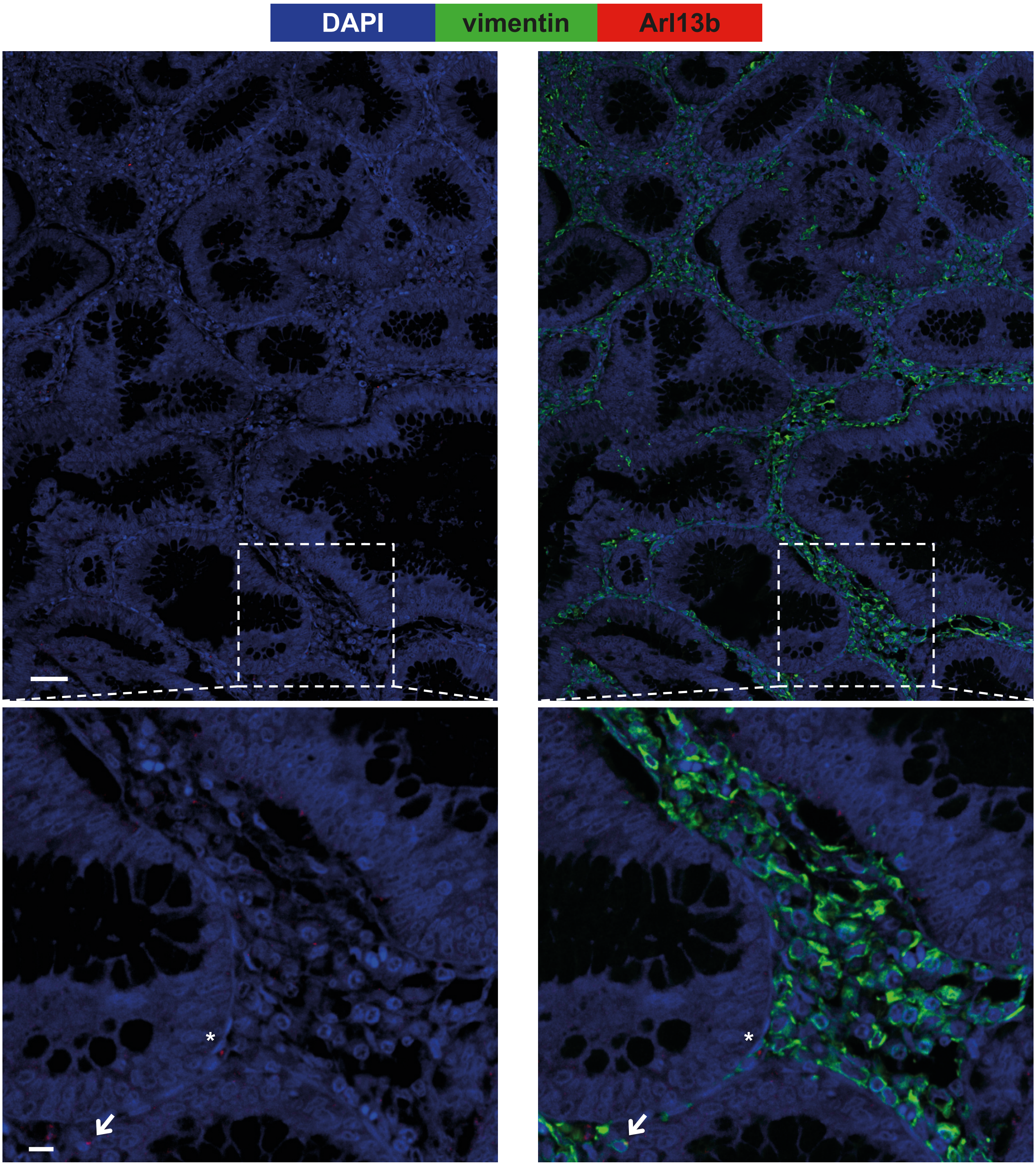

**Supplementary figure 7B**

**Peritumoral tissue (Stage 1)**

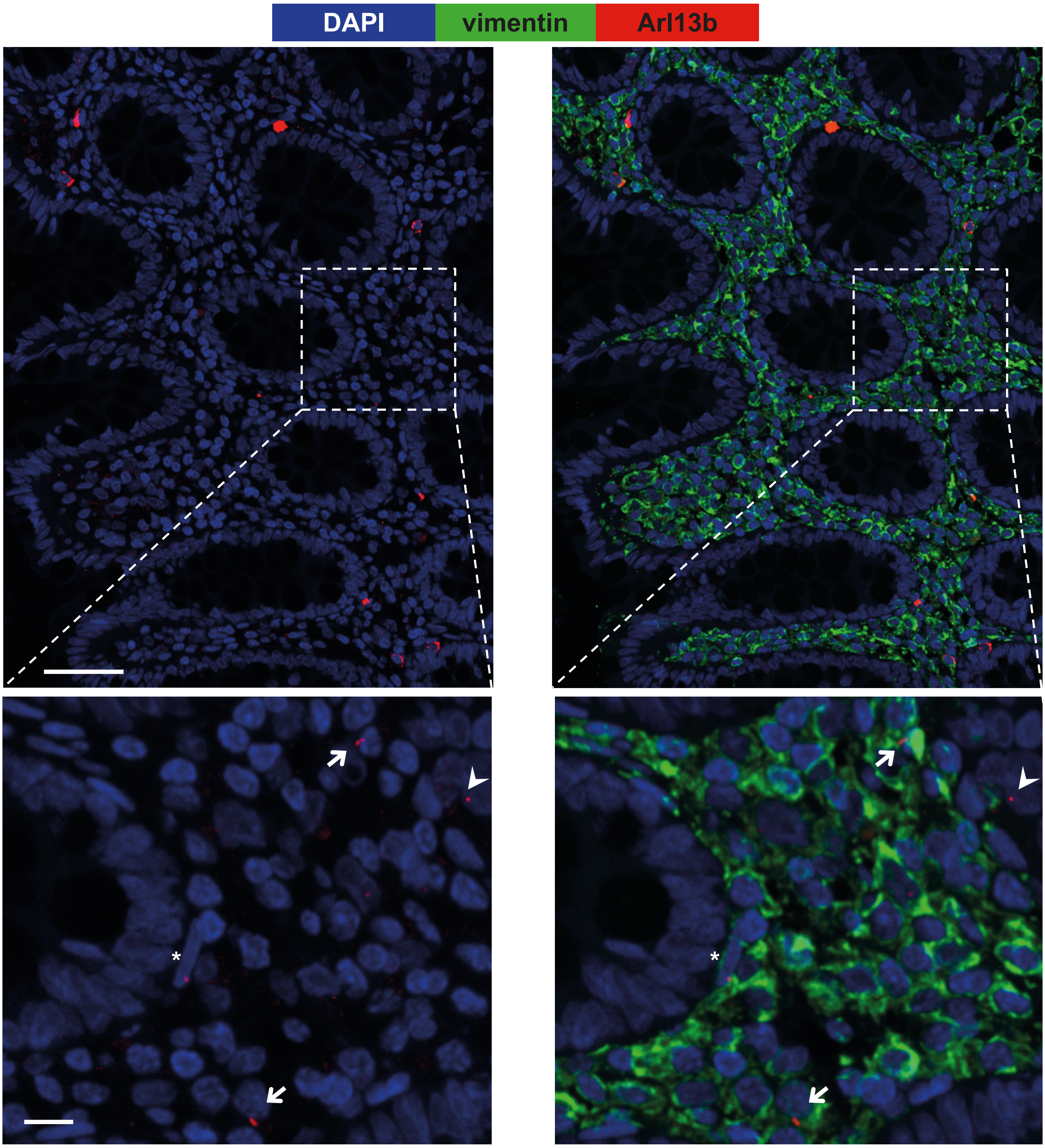
